# SCassist: An AI Based Workflow Assistant for Single-Cell Analysis

**DOI:** 10.1101/2025.04.22.650107

**Authors:** Vijayaraj Nagarajan, Guangpu Shi, Samyuktha Arunkumar, Chunhong Liu, Jaanam Gopalakrishnan, Pulak R Nath, Junseok Jang, Rachel R Caspi

## Abstract

**Summary:** Single-cell RNA sequencing (scRNA-seq) data analysis often involves complex iterative workflow, requiring significant expertise and time. To navigate this complexity, we have developed SCassist, an R package that leverages the power of the large language models (LLM’s) to guide and enhance scRNA-seq analysis. SCassist integrates LLM’s into key workflow steps, to analyze user data and provide relevant recommendations for filtering, normalization and clustering parameters. It also provides LLM guided insightful interpretations of variable features and principal components, along with cell type annotations and enrichment analysis. SCassist provides intelligent assistance using popular LLM’s like Google’s Gemini, OpenAI’s GPT and Meta’s Llama3, making scRNA-seq analysis accessible to researchers at all levels.

**Availability and implementation:** The SCassist package, along with the detailed tutorials, is available at GitHub. https://github.com/NIH-NEI/SCassist

## Introduction

The standard Single-cell analysis workflow includes several steps that require researchers to identify and use appropriate parameters. These steps include quality filtering, normalization, dimension reduction and clustering (Nath et al. 2024) . Other aspects, such as understanding the context of variable genes, principal components, cell type markers and enrichment analysis, require extensive knowledge of the background biology to extract meaningful insights from the data.

Large Language Models (LLM’s) have significantly advanced text generation, text summarization, translation, question answering, image generation, audio/video generation, code generation etc., (Tom B. Brown 2020). In biomedical research and clinical settings, LLM based applications are being explored for their potential in drug design, drug discovery, sequence analysis, target discovery, clinical diagnosis, treatment recommendations, and outcome predictions (Ji et al. 2021; Lee et al. 2020; Thirunavukarasu et al. 2023). Researchers have also integrated LLM-generated results into radiology reporting workflows (Jorg et al. 2024). Although ‘hallucination’ is recognized as an inherent limitation of LLM’s (Kankanhalli 2024), new methods and tools to mitigate this and take advantage of the LLM’s abilities, like retrieval augmentation, are also being established (Jin et al. 2024) .

In the field of single-cell analysis, various tools and methods have been developed using specialized, foundation or fine-tuned models to address challenges such as cell type annotation, imputation, integration, clustering, dimensionality reduction, and trajectory analysis (Szalata et al. 2024). Each of these current AI based single-cell analysis tools come with distinct strengths. Models like Geneformer (Theodoris et al. 2023), scGPT (Cui et al. 2024), scBERT (F. Yang, Wang, W., Wang, F. et al. 2022), TOSICA (Chen et al. 2023), CellLM (Nie 2023), GeneCompass (X. Yang et al. 2024) and CellPLM (Hongzhi Wen 2023) are primarily specialized towards identifying and annotating cell populations, while models like scTPC (Qiu et al. 2024), tGPT (Shen et al. 2023) are geared towards advanced approaches for cell clustering and lineage identification. In addition to the cell clustering tasks or cell annotation tasks, Geneformer, scGPT, CellLM, scFoundation (Hao et al. 2024), GeneCompass and CellPLM are also able to handle supervised or zero-shot tasks like perturbation effect prediction and drug response prediction.

Apart from the direct use of single-cell trained models for carrying out specialized tasks, several applications are also built on top of these models, in an effort to leverage their potential use in understanding the single-cell data. For example, GPTCelltype (Hou and Ji 2024) leverages the impressive reasoning capabilities of GPT-4 to automate cell type annotation with high accuracy, offering a streamlined solution for this crucial task. ChatCell (Fang 2024) introduces a novel natural language interface, enabling users to interact with scRNA-seq data through intuitive commands and perform specialized tasks like cell generation to drug sensitivity prediction, showcasing the versatility of fine-tuned language models.

However, while these tools offer significant progress in specific areas, SCassist uniquely distinguishes itself by aiming to provide comprehensive workflow guidance across the entire scRNA-seq analysis pipeline, a feature absent in these task-specific tools. SCassist aims to empower users with step-by-step recommendations and insightful interpretations at each stage (Figure 1), fostering deeper understanding and control within the familiar R/Seurat environment. Furthermore, SCassist's novelty lies in its pioneering workflow assistance paradigm that leverages the accessibility and evolving power of general-purpose Large Language Models, offering a more flexible, cost-effective, and user-centric approach compared to specialized models or annotation-centric solutions. This holistic approach positions SCassist not just as another tool, but as a novel and complementary intelligent assistant that aims to enhance the entire scRNA-seq analysis journey. A more detailed feature comparison is provided in Supplementary Table 1, to highlight novel contributions of SCassist, compared to selected existing methods.

**Figure 1:**
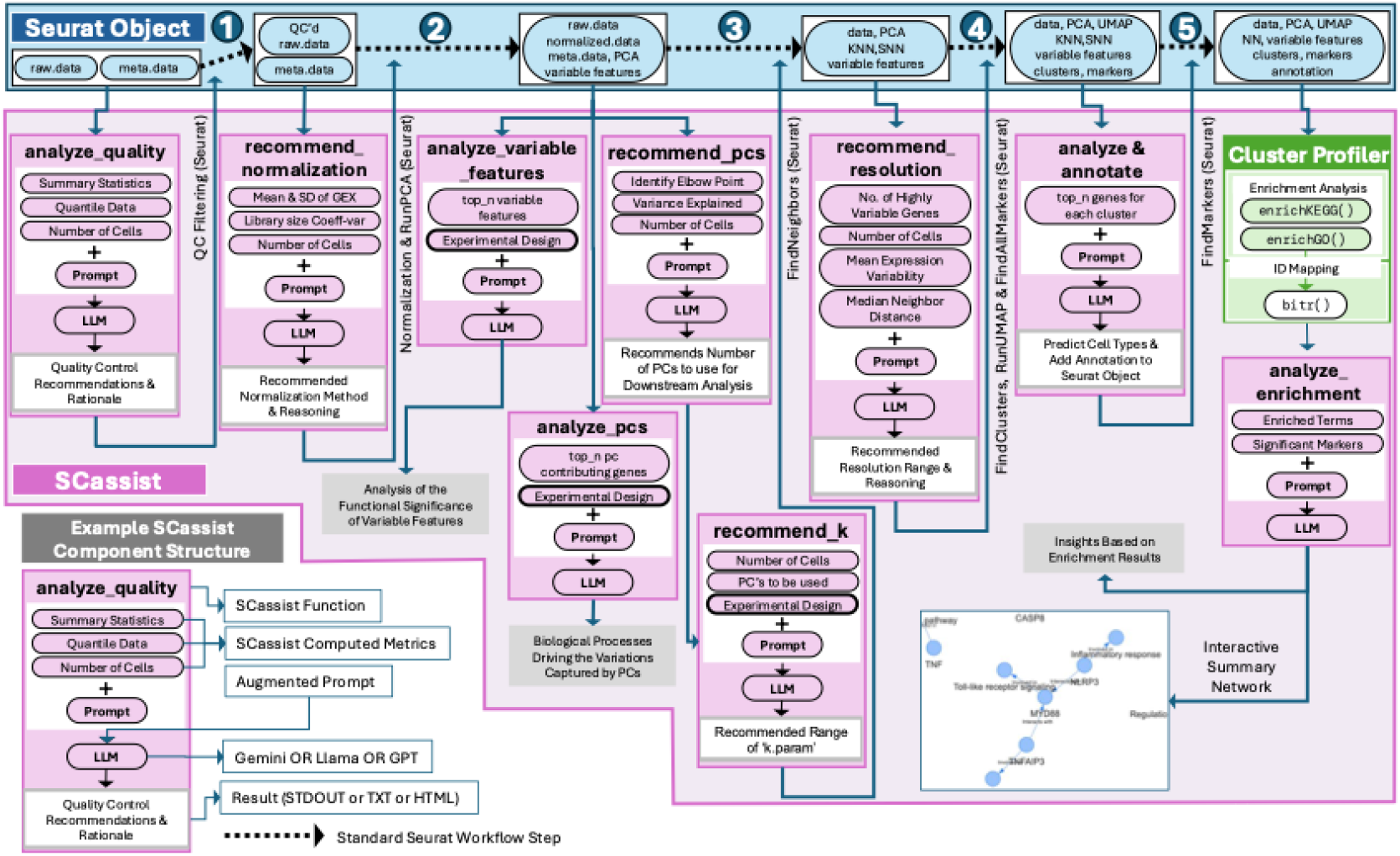
The general architecture of the SCassist algorithm. SCassist, an LLM-powered assistant, streamlines single-cell analysis within the standard Seurat workflow. The top portion of the figure depicts the typical Seurat steps (quality control, normalization, dimensionality reduction, clustering, and annotation), while the interconnected pink boxes represent SCassist components, providing data-driven insights and parameter recommendations for each step. SCassist could be used at any stage of the standard single-cell workflow, starting from the quality control stage, where the user input for SCassist is simply the Seurat object containing the raw count matrix data. For the given Seurat object, SCassist generates metrics like summary statistics, quantile data, variance explained, and others. These metrics are then used to build augmented prompts for large language models (LLMs), recommending optimal parameters for filtering, normalization, dimensionality reduction, identifying significant features and offering insights (from variable genes, principal components, differentially expressed genes), and annotating clusters along with detailed reasoning.

As an opensource R package, SCassist helps the researchers easily navigate through the complex and time-consuming components of the single-cell transcriptomics data analysis. SCassist leverages the retrieval and tool-based augmentation techniques to seamlessly integrate the power of the LLM’s into the standard single-cell analysis workflow. This enables data exploration and understanding through insightful descriptions and enhanced interpretability of complex concepts, accelerating discovery.

## Methods

### Architecture

SCassist is written in the R programming language. SCassist takes the Seurat single-cell object and related Seurat results to generate relevant data/metrics. Seurat results are also used to perform enrichment analysis with ClusterProfiler, generating additional tools-based data/metrics. An augmented prompt is then constructed, by appropriately combining the data/metrics and experiment description using the prompt template. The contextualized augmented prompt is then submitted to the LLM. The LLM’s response is parsed, and the results are displayed on the screen and/or saved them to a file, or appended to the Seurat object, as metadata. In some functions, the parsed response from the LLM is also used as the input for SCassist’s second round of LLM queries. The general architecture of SCassist is illustrated in Figure 1. The SCassist package was built using R 4.4.1 running on the macOS Ventura platform.

### Large Language Models

SCassist allows users to choose between “Google”, “OpenAI” or “Ollama” as the LLM server. The default models are set as “gemini-1.5-flash-latest” for Google, “gpt-4o-mini” for OpenAI and “llama3” for Ollama. If using Google or OpenAI as the server, the user is expected to provide an API key from Google or OpenAI. For Ollama, the user is expected to have it installed and running locally. We chose Google as a commercial online LLM server, due to their large context window availability. Ollama was chosen as an open-source, offline LLM server, providing free access to open-source models like Llama3 locally, without sending the data to a remote server. Detailed installation and setup instructions are available on our project’s <u>GitHub page</u>.

### Implementation

The SCassist package implements several key functions as described below:

- **SCassist_analyze_quality():** Analyzes the quality of the single-cell data using metrics like nCount_RNA, nFeature_RNA, and any user-specified quality metrics generated through Seurat’s PercentageFeatureSet, to recommend filtering cutoffs.
- **SCassist_recommend_normalization():** Analyzes the characteristics of the dataset (number of cells, gene expression distribution, library size variation) to recommend the most suitable normalization method from Seurat's options.
- **SCassist_analyze_variable_features():** Analyzes the top variable features identified by Seurat to identify enriched gene ontologies or pathways, based on known feature characteristics and explain their relevance to the experimental design.
- **SCassist_recommend_pcs():** Analyzes the variance explained by each principal component (PC) to recommend the optimal number of PCs for downstream analysis.
- **SCassist_analyze_pcs():** Analyzes the top PCs to understand the driving biological processes based on top contributing genes and their associated pathways.
- **SCassist_recommend_k():** Analyzes the number of cells, PCs used, and clustering goals to recommend a range of potential k.param values for FindNeighbors
- **SCassist_recommend_res():** Analyzes the number of cells, mean expression variability, and median neighbor distance to recommend a suitable resolution range for FindClusters.
- **SCassist_analyze_and_annotate():** Analyzes top markers for each cluster to predict potential cell types based on the markers and provide reasoning. Optionally annotates the Seurat object with the predicted cell types.
- **SCassist_analyze_enrichment():** Runs ClusterProfiler for KEGG pathway and Gene Ontology enrichment. Analyzes and processes the ClusterProfiler results to integrate and generate insights on significant pathways, potential regulators, key genes or targets, and a summary network.
- **SCassist_summary_network():** Uses the LLM to extract network data from the output of SCassist_analyze_enrichment and creates an interactive network visualization that summarizes the system.

A detailed overview of all the SCassist functions, including their specific tasks, key benefits and the typical Seurat workflow stage at which they are applied, are presented in the Supplementary Table 2.

Detailed documentation for each of the functions and methods, along with usage and examples, are provided with corresponding R help files, as part of the SCassist package. Each of the above steps can be run in sequence, along with the appropriate standard Seurat single-cell analysis workflow steps. Each of the above steps can also be run independently at any appropriate step of the standard workflow. SCassist also provides options for advanced users to choose specific models and set custom model parameters.

### Prompt Integration within SCassist Functions

Here we detail how template prompts are utilized within the SCassist functions. Users do not directly specify prompts as input parameters to the individual SCassist functions (e.g., `SCassist_analyze_quality(),`

`SCassist_recommend_normalization()`). Instead, each function is designed to internally construct an augmented prompt based on the input data (typically a Seurat object or results from previous steps), computed metrics (e.g., mean expression variability, median neighbor distance, quantile data etc.,) and a pre-defined prompt template. These prompt templates (Supplementary File 1), engineered to elicit specific insights from the LLM, are integral to each function’s operation.

For example, as illustrated in Figure 1, `SCassist_analyze_quality()` takes a Seurat object with raw single-cell data and its corresponding metadata (nCount_RNA, nFeature_RNA, etc.,) as input, calculates relevant data metrics (Summary statistics, Quantile data and Number of cells), and then uses these metrics to populate a pre-defined prompt template, to create an augmented prompt that is sent to the LLM to recommend filtering cutoffs. The recommended filtering cut-off values, along with relevant reasoning from the LLM, are displayed to the user. The user then uses these parameters, as a guidance, to move on to the next step, which is quality filtering. The actual prompt template for `SCassist_analyze_quality()` and an example output using that template is provided in Supplementary File 2.

The prompt templates themselves are designed to be modular and adaptable, allowing for future refinement and customization. The full set of prompt templates used by each function is available on our project’s <u>GitHub page</u>, providing transparency and enabling advanced users to modify the prompts if desired. This approach ensures that users can leverage the power of LLMs without needing to manually craft complex prompts, streamlining the analysis process and promoting ease of use. Depending on the function used, either the direct response or the parsed response from the LLM is then integrated back into the function’s output, providing data-driven recommendations and insightful interpretations.

The SCassist R package can be installed and run on any operating system, using R versions 4.4.1 and above. The R packages Seurat, rollama, httr, jsonlite, visNetwork, clusterProfiler and BiocManager are required for the full functioning of SCassist.

## Evaluation

### Dataset

To evaluate SCassist’s performance, we used datasets from two of our recently published single-cell transcriptome studies (Supplementary Table 3). One study identified an NK cell subset associated with disease activity in human Uveitis patients (Nath et al. 2024) and the other identified a CTCF binding motif site critical for mouse Th1 cell fate specification (Liu et al. 2024). We reanalyzed these datasets using SCassist and evaluated the SCassist-generated workflow reports against the previously published standard workflow reports for these two datasets.

### Groundedness score

SCassist’s ability to use the provided content was measured using the groundedness score. This score serves as an indirect measure of hallucination by estimating how well the LLM’s response is grounded in the provided information. As the ground truth, we used all the data-derived metrics, lists of variable genes, lists of genes from PCs, lists of genes from cluster markers, lists of genes from ClusterProfiler enrichment results, and lists of genes derived from SCassist’s enrichment analysis summary reports. We used GeneVenn (Pirooznia et al. 2007) to compare the terms in the SCassist’s responses with the corresponding terms in our ground truth, to score for the overlaps. The groundedness score (G) was calculated as the cardinality of the intersection between the ground truth (GT) and the LLM’s response (LLM), divided by the cardinality of the LLM’s response: G = |GT ∩ LLM / |LLM|.

### Contextual relevance

The contextual relevance of the LLM’s response in SCassist was measured using semantic similarity. We compared the SCassist’ enrichment analysis summary response to that of the input document containing enriched pathways and GO terms, using the transformer-based language model BERT (Bidirectional encoder representations from transformers) (Devlin et al. 2018). Briefly, the contents were tokenized, added with relevant special tokens to mark the start and end of the contents and chunked to fit BERT’s input length. We then converted the chunks into input ids and extracted relevant embeddings. The chunked embeddings were combined and used to compute the cosine similarity. The semantic similarity analysis was performed in a python virtual environment using the transformers, torch and sklearn.metrics.pairwise packages. The fully documented python script used for this analysis is provided in our project’s GitHub page.

### Human Evaluation

We assessed the quality of the SCassist generated workflow reports against the standard single-cell analysis workflow reports using the Likert scale (1 to 5), to account for Accuracy, Relevance, Clarity, Trustworthiness and Overall satisfaction (Likert 1932). We designed our scoring sheet with clear anchors, describing how to differentiate the scores, including specific descriptions to reduce any ambiguity. We chose four senior level scientists (two Senior Scientist’s from BioTech industry and two Staff Scientist’s from the NIH) and four junior level scientists (Post Doctoral Fellows and Post Baccalaureate Fellows from the NIH) to evaluate our reports. Two of the senior scientists were the original authors of the published workflows that are analyzed with SCassist in this study. The template scoring sheet is provided in Supplementary Table 4.

### Statistics

For the expert human evaluation, we performed the Wilcoxon Signed-Rank test to assess whether the overall Likert scale scores were significantly skewed towards higher values, indicating positive user perception of the LLM's performance. We also performed the Wilcoxon Rank-Sum test to evaluate if the Likert scale scores are affected by the level of expertise and by the dataset evaluated. Summary statistics, along with frequency distribution plots were used to understand the expert human evaluation scores.

## Results

SCassist is available as an opensource R package at our project GitHub page: https://github.com/NIH-NEI/SCassist. Detailed tutorials, walking through the step-by-step workflows, using example data, are also provided at the GitHub project site. SCassist package could be installed using the R command: `devtools::install_github(“NIH-NEI/SCassist”)`. Detailed tutorials describing the usage of commands and corresponding responses are shared through the project GitHub page.

Recommendation Process

To enhance the user’s understanding and ensure the alignment of recommendations with their analytical needs, SCassist explicitly details the decision-making process as part of each functions output. For instance, the below example output (part of the SCassist Tutorial) for `SCassist_recommend_res()` demonstrates how these recommendations are derived. The user calls the function as below, with the Seurat object as the input (`allsamplesgood,`before the standard “Clustering” step in their single-cell analysis workflow;

~~~
SCassist_recommend_res(“allsamplesgood", llm_server="google",
api_key_file = api_key_file)
~~~

SCassist, analyzes the Seurat object, computes key dataset characteristics (such as mean expression variability, median neighbor distance, number of highly variable genes and total number of cells), combines it with the corresponding prompt template, runs it through the LLM and provides the data-driven recommendations, as shown in the below example output;

~~~
## Based on the data characteristics, I recommend:
## **Recommended Resolution:** seq(0.2, 1.2, 0.1)
##
## **Reasoning:**
## The mean expression variability of 0.868717336264306 suggests a moderate level of heterogeneity in your dataset. This indicates that there are likely distinct cell populations, but they may not be drastically different from each other. The median neighbor distance of 2.50502443313599 in the k-nearest neighbor graph further supports this notion, as it implies a moderate level of separation between cells.
## Therefore, a resolution range of 0.2 to 1.2 with increments of 0.1 is recommended. This range allows for the identification of both distinct and subtle cell populations. Lower resolutions (closer to 0.2) will capture broader, more general cell types, while higher resolutions (closer to 1.2) will reveal finer distinctions within those populations. By exploring this range, you can effectively identify the optimal resolution for your specific analysis goals.
~~~

The above result provides not only the recommended resolution range (e.g., seq(0.2, 1.2, 0.1)) but also the reasoning behind the recommendation. This reasoning explicitly states the criteria used (e.g., “The mean expression variability of 0.868717336264306 suggests a moderate level of heterogeneity…”). The inclusion of this reasoning clarifies which aspects of the data the LLM has considered and how these data points influence the function's suggestions, allowing users to assess the relevance of the recommendations in relation to their specific analysis goals and to determine whether the criteria used align with their analytic needs. By making the underlying logic transparent, we aim to build trust in SCassist’s recommendations and empower users to make informed decisions about their scRNA-seq analysis.

To assess the accuracy of cell type annotations performed by `SCassist_analyze_and_annotate(),` we compared its results to those obtained using GPTCelltype, using the same FindAllMarkers results as input (generated from the example data - GSM6625298, provided in SCassist’s GitHub page). The resulting cell type assignments, presented in Supplementary File 3, were highly concordant. Importantly, beyond the cell type labels, `SCassist_analyze_and_annotate()` provides clear and transparent reasoning for each annotation call, as is consistent throughout all of SCassist's functions. This enables users to easily evaluate and understand the basis for the annotation.

## Evaluation

### Groundedness score

Using a total of 4136 tokens as ground truth, for different categories, SCassist scored an average of 98.7% groundedness for the LCMV. For the BCRUV data, using a total of 4110 tokens, SCassist scored an impressive average of 99.9% groundedness. Detailed scores for different categories are presented in the Supplementary Table 5.

### Semantic similarity

Using BERT’s uncased model, SCassist scored 76% on the semantic similarity between the terms used and the reports generated, for the LCMV data. SCassist scored a similar 74% for the BCRUV data. Detailed scores for different categories are presented in the Supplementary Table 6.

### Expert human evaluation

Evaluation of SCassist’s workflow reports, against the standard workflow reports, by eight human expert evaluators with different levels of expertise show that SCassist has good accuracy and generally performs well on relevance and trustworthiness. SCassist performs exceptionally well on clarity, with most evaluators (7 out of 8) finding it useful (Supplementary Figure 1). Detailed scores are provided in Supplementary Table 7. The Wilcoxon Signed-Rank test with a p-value of 0.0001122 revealed a statistically significant skew towards higher scores, indicating that SCassist performs well overall. The Wilcoxon Rank-Sum test revealed no statistically significant differences in the LLM's performance across expert levels and across the two different datasets (Supplementary Table 8).

### Cost

We made a total of 1,978 API calls to Google Gemini, in the two-month testing period during this project. The total cost for the 1,978 API calls was $2.07 excluding taxes. Running SCassist using the local Ollama server option did not incur any direct costs.

### Limitations

Due to the fast-changing landscape of the LLM’s, SCassist’s responses in the future cannot be predicted, due to possible changes in the underlying models. That said, we believe the responses would only get better. We commit to monitoring the test workflow for any substantial changes in the responses at a minimum once every three months and to provide appropriate and diligent updates to the SCassist package. We will strive to fix any issues reported by the users, at the earliest possible time.

## Conclusion

SCassist consistently achieved high scores across groundedness, semantic similarity and expert human evaluation, indicating its ability to generate accurate and meaningful insights. Our analysis also revealed that SCassist’s performance is robust across different datasets and the evaluation scores are not affected by differences in the expertise level of the evaluator. The user-friendly interface and integration with popular LLM’s make SCassist accessible to researchers at all levels. It empowers them to leverage AI-driven insights to accelerate their single-cell analysis workflow, while at the same time uncovering deeper biological insights and advancing scientific discovery. Our future development efforts will extend SCassist's capabilities to support multi-modal and spatial workflows, including interaction and trajectory analyses. We also plan to integrate the LLMs' function-calling abilities to offer automation.

## Data availability

All data and code discussed in this work are available in our GitHub page at : https://github.com/NIH-NEI/SCassist/. Snapshot of the code and documentation is also available at https://doi.org/10.5281/zenodo.15298665.

## Acknowledgments

This work utilized the computational resources of the NIH HPC Biowulf cluster. The authors sincerely thank Drs. Charles Egwuagu and Han-Yu Shih for their support and guidance throughout the related historical work. The authors also thank Zixuan Peng for help with evaluation and review of the manuscript. We also thank the NEI IRP program for funding this research.

## Supplementary information

SCassist_Supplementary_Data.pdf

SCassist_Supplementary_File_1.pdf

SCassist_Supplementary_File_2.pdf

SCassist_Supplementary_File_3.pdf

SCassist_Supplementary_Table_1.xlsx

SCassist_Supplementary_Table_2.xlsx

## References

Chen, J., et al. (2023), ‘Transformer for one stop interpretable cell type annotation’, Nat Commun, 14 (1), 223.

Cui, H., et al. (2024), ‘scGPT: toward building a foundation model for single-cell multi-omics using generative AI’, Nat Methods, 21 (8), 1470–80.

Devlin, Jacob, et al. (2018), ‘BERT: Pre-training of Deep Bidirectional Transformers for Language Understanding’, arxiv.org.

Fang, Yin and Liu, Kangwei and Zhang, Ningyu and Deng, Xinle and Yang, Penghui and Chen, Zhuo and Tang, Xiangru and Gerstein, Mark and Fan, Xiaohui and Chen, Huajun ‘ChatCell: Facilitating Single-Cell Analysis with Natural Language’.

Hao, M., et al. (2024), ‘Large-scale foundation model on single-cell transcriptomics’, Nat Methods, 21 (8), 1481–91.

Hongzhi Wen, Wenzhuo Tang, Xinnan Dai, Jiayuan Ding, Wei Jin, Yuying Xie, Jiliang Tang (2023), ‘CellPLM: Pre-training of Cell Language Model Beyond Single Cells’, bioRxiv.

Hou, W. and Ji, Z. (2024), ‘Assessing GPT-4 for cell type annotation in single-cell RNA-seq analysis’, Nat Methods, 21 (8), 1462–65.

Ji, Y., et al. (2021), ‘DNABERT: pre-trained Bidirectional Encoder Representations from Transformers model for DNA-language in genome’, Bioinformatics, 37 (15), 2112–20.

Jin, Q., et al. (2024), ‘GeneGPT: augmenting large language models with domain tools for improved access to biomedical information’, Bioinformatics, 40 (2).

Jorg, T., et al. (2024), ‘A novel reporting workflow for automated integration of artificial intelligence results into structured radiology reports’, Insights Imaging, 15 (1), 80.

Kankanhalli, Ziwei Xu and Sanjay Jain and Mohan (2024), ‘Hallucination is Inevitable: An Innate Limitation of Large Language Models’, arXiv.

Lee, J., et al. (2020), ‘BioBERT: a pre-trained biomedical language representation model for biomedical text mining’, Bioinformatics, 36 (4), 1234–40.

Likert, Rensis (1932), ‘A Technique for the Measurement of Attitudes’, Archives of Psychology, 140, 1–55.

Liu, C., et al. (2024), ‘A CTCF-binding site in the Mdm1-Il22-Ifng locus shapes cytokine expression profiles and plays a critical role in early Th1 cell fate specification’, Immunity, 57 (5), 1005–18 e7.

Nath, P. R., et al. (2024), ‘Single-cell profiling identifies a CD8(bright) CD244(bright) Natural Killer cell subset that reflects disease activity in HLA-A29-positive birdshot chorioretinopathy’, Nat Commun, 15 (1), 6443.

Nie, Suyuan Zhao and Jiahuan Zhang and Zaiqing (2023), ‘Large-Scale Cell Representation Learning via Divide-and-Conquer Contrastive Learning’, arXiv.

Pirooznia, M., Nagarajan, V., and Deng, Y. (2007), ‘GeneVenn - A web application for comparing gene lists using Venn diagrams’, Bioinformation, 1 (10), 420–2.

Qiu, Y., et al. (2024), ‘scTPC: a novel semisupervised deep clustering model for scRNA-seq data’, Bioinformatics, 40 (5).

Shen, H., et al. (2023), ‘Generative pretraining from large-scale transcriptomes for single-cell deciphering’, iScience, 26 (5), 106536.

Szalata, A., et al. (2024), ‘Transformers in single-cell omics: a review and new perspectives’, Nat Methods, 21 (8), 1430–43.

Theodoris, C. V., et al. (2023), ‘Transfer learning enables predictions in network biology’, Nature, 618 (7965), 616–24.

Thirunavukarasu, A. J., et al. (2023), ‘Large language models in medicine’, Nat Med, 29 (8), 1930–40.

Tom B. Brown, Benjamin Mann, Nick Ryder, Melanie Subbiah, Jared Kaplan, Prafulla Dhariwal, Arvind Neelakantan, Pranav Shyam, Girish Sastry, Amanda Askell, Sandhini Agarwal, Ariel Herbert-Voss, Gretchen Krueger, Tom Henighan, Rewon Child, Aditya Ramesh, Daniel M. Ziegler, Jeffrey Wu, Clemens Winter, Christopher Hesse, Mark Chen, Eric Sigler, Mateusz Litwin, Scott Gray, Benjamin Chess, Jack Clark, Christopher Berner, Sam McCandlish, Alec Radford, Ilya Sutskever, and Dario Amodei (2020), ‘Language models are few-shot learners’, NIPS’20: Proceedings of the 34th International Conference on Neural Information Processing Systems (159; Vancouver, BC, Canada), 1877 - 901.

Yang, F., Wang, W., Wang, F. et al. (2022), ‘scBERT as a large-scale pretrained deep language model for cell type annotation of single-cell RNA-seq data’, Nat Mach Intell, 4, 852–66.

Yang, X., et al. (2024), ‘GeneCompass: deciphering universal gene regulatory mechanisms with a knowledge-informed cross-species foundation model’, Cell Res, 34 (12), 830–45.

