## Supplementary File 1 for "SCassist: An AI Based Workflow Assistant for Single-Cell Analysis"

**Supplementary File 1:** Overview of prompt templates used in SCassist functions.

### SCassist R Package Prompts Report

This report outlines the prompts used in the SCassist R package functions for interacting with large language models (LLMs).

**Documentation by :** Gemini \
**Documentation reviewed by :** Vijay Nagarajan PhD

**1. SCassist_analyze_quality**

```
  I have summary statistics and quantile data for nCount_RNA, nFeature_RNA, [quality_metrics], from a single cell experiment.
  I want to determine refined cutoff values that are more sensitive to the tails of each distribution, by combining both the summary statistics and the quantile information. The goal is to filter out poor quality cells.

  **nCount_RNA**
  **Summary Statistics:**
  * Minimum: [nCount_summary["Min."]]
  * 1st Quartile: [nCount_summary["1st Qu."]]
  * Median: [nCount_summary["Median"]]
  * Maximum: [nCount_summary["Max."]]

  **Quantile Data:**
  * 5th Percentile: [nCount_quantiles["5%"]]
  * 95th Percentile: [nCount_quantiles["95%"]]

  **nFeature_RNA**
  **Summary Statistics:**
  * Minimum: [nFeature_summary["Min."]]
  * 1st Quartile: [nFeature_summary["1st Qu."]]
  * Median: [nFeature_summary["Median"]]
  * Maximum: [nFeature_summary["Max."]]

  **Quantile Data:**
  * 5th Percentile: [nFeature_quantiles["5%"]]
  * 95th Percentile: [nFeature_quantiles["95%"]]

  [Summary statistics for each quality metric]

  ** Total number of cells : ** [nCells]

  **Please provide the refined lower and upper cutoff values for nCount_RNA and nFeature_RNA, [and the upper cutoff values for [quality_metrics]], calculated based on the provided data and taking into account the potential presence of tails in the distribution. Dont generate the cuffoff ONLY based on the percentiles, instead combine the percentiles with a logic that we need to remove outliers, but still keep good data. Make sure to state that the researcher should test a range of values around your recommendation**. Do not provide any IMPORTANT NOTES. Start the response saying that, based on the your data summary, below are my recommendations for the quality filtering of the data
```

**2. SCassist_recommend_normalization**

  ```
  Consider the specific characteristics of the single-cell RNA-seq dataset, such as the number of cells, the distribution of gene expression values, and the observed library size variations. Recommend the most appropriate normalization method for this dataset from the following options available in Seurat: [seurat_normalization_methods]. Explain why this method is preferred and discuss any potential alternatives.)

  The single-cell RNA-seq dataset contains [num_cells] cells.  The mean gene expression is [gene_expr_mean] with a standard deviation of [gene_expr_sd]. The coefficient of variation for library sizes is [library_size_cv].
  ```

**3. SCassist_analyze_variable_features**

  ```
  Analyze the following list of genes: [gene_list]
  These genes were identified as variable features. Identify the most enriched gene ontologies or pathways among these genes. Dont provide any pvalues or enrichment scores, just summarize based on these genes known functions. Dont provide any Further Analysis suggestions
  [If experimental_design is provided, add: Explain the relevance of these categories to the experimental design: [experimental_design].]
  ```

**4. SCassist_recommend_pcs**

  ```
  [If experimental_design is provided, add: I have a single-cell experiment where I performed principal component analysis (PCA) on [experimental_design].]
  I have a single-cell experiment where I performed principal component analysis (PCA). The variance explained by each PC is: [variance_text]
  Based on this information, determine the optimal number of PCs to use for downstream analysis, such as finding neighbors or running UMAP. Explain your reasoning and consider the following factors:

  * **Elbow point:** Is there a clear 'elbow' in the scree plot?
  * **Variance explained:**  What percentage of the total variance is captured by the chosen number of PCs?
  * **Balance between complexity and interpretability:** A higher number of PCs might capture more subtle variation but make the analysis more complex.

  Provide your recommendation as a single number (e.g., 5) and a concise explanation.
  ```

**5. SCassist_analyze_pcs**

  ```
  [experimental_design]
  We performed QC, normalization and PCA on this data using Seurat. Here is the list of top PC's and their genes from our analysis:
  ```

  PC[i]: [genes]

  Identify the top contributing genes for this PC. Based on the gene functions and biological pathways associated with these genes, suggest potential biological processes that might be driving the variations captured by this PC. Present your results in a short paragraph

  [Combined prompt for overall summary based on individual PC summaries]
  ```

**6. SCassist_recommend_k**

  ```
  I'm analyzing single-cell RNA sequencing data from [experimental_design]
  The dataset contains approximately [num_cells] cells.
  I've determined that using [num_pcs] PCs (`dims` = [num_pcs]) is suitable for my data.
  I'm interested in identifying distinct cell populations.
  My goal is to [clustering_goal].
  Can you suggest a range of potential `k.param` values, in whole number, to explore for `FindNeighbors()` based on this information? Provide the output as, Recommended K: and a two short reasoning paragraph under Reasoning:
  ```

**7. SCassist_recommend_res**

  ```
  I am analyzing single-cell RNA sequencing data. My dataset consists of [data_for_llm$n_cells] cells and [data_for_llm$n_genes] genes.
  The mean expression variability across genes is [data_for_llm$mean_expression_variability]
  and the median neighbor distance in the k-nearest neighbor graph is [data_for_llm$median_neighbor_distance].

  What resolution range would be most suitable for identifying distinct and subtle populations of cells in my data?
  Provide the output as, Recommended Resolution, EXAMPLE: seq(starting resolution number,ending resolution number,increment number), : and a short reasoning paragraph under Reasoning. Start your response saying that, based on the data characterisitcs i recommend;
  ```

**8. SCassist_analyze_and_annotate**

  ```
  The provided genes are the top markers of this single cell cluster. Analyze it and predict a potential cell type based on the markers. provide output in three columns. the first column should be the cluster number, second column should be the name of the potential cell type, third column should be a one paragraph reasoning. separate the columns using a colon. dont provide any other additional content generated by you, like: here is the analysis, etc. Here is an EXAMPLE output - '8:Megakaryocyte-precursor cells: The combination of markers LY6G6F, GP9, ITGA2B, and TMEM40 suggests a megakaryocytic origin, with involvement in platelet development and function'. Here is the input cluster number and corresponding markers for your analysis:
  cluster [cluster_num]: [cluster_markers]
  ```
  ```

**9. SCassist_analyze_enrichment**

```
This below Data is a list of [KEGG/GO] pathway enrichment results for a set of differentially expressed genes obtained from a single cell experiment involving [experimental_design].

Data:
[json_enrichment]

Analyze all of the pathways/concepts from my above Data and provide insights in a structured format. Include:

1. **Significant Pathways/Concepts:** Analyze all the pathways/concepts in my list, in the context of the system involving [experimental_design], and summarize common themes, in 2 paragraph, with a total of 10 lines.
2. **Regulators:** Include potential involvment of any transcription factors from the given list, in the context of the system involving [experimental_design], in a 5 line paragraph.
3. **Key Genes or Potential Targets:** Suggest which genes from the enriched pathways/ontologies, in my data input, might be important to the system or a target based on their potential impact on the system involving [experimental_design], in a 10 line paragraph.

Do not provide any Further Investigation or comments, not asked for.

[Prompt for extracting network data from the LLM response]
```

This report provides a comprehensive overview of the prompts used in the SCassist R package. These prompts are designed to guide the LLM in providing insightful and relevant information for single-cell RNA sequencing data analysis.
