## Supplementary File 2 for "SCassist: An AI Based Workflow Assistant for Single-Cell Analysis"

**Supplementary File 2:** Prompt Template and Example Output for `SCassist_analyze_quality()` function.

Prompt Template;

```
  ```
  I have summary statistics and quantile data for nCount_RNA, nFeature_RNA,
[quality_metrics], from a single cell experiment.
  I want to determine refined cutoff values that are more sensitive to the
tails of each distribution, by combining both the summary statistics and the
quantile information. The goal is to filter out poor quality cells.

  **nCount_RNA**
  **Summary Statistics:**
  * Minimum: [nCount_summary["Min."]]
  * 1st Quartile: [nCount_summary["1st Qu."]]
  * Median: [nCount_summary["Median"]]
  * Maximum: [nCount_summary["Max."]]

  **Quantile Data:**
  * 5th Percentile: [nCount_quantiles["5%"]]
  * 95th Percentile: [nCount_quantiles["95%"]]

  **nFeature_RNA**
  **Summary Statistics:**
  * Minimum: [nFeature_summary["Min."]]
  * 1st Quartile: [nFeature_summary["1st Qu."]]
  * Median: [nFeature_summary["Median"]]
  * Maximum: [nFeature_summary["Max."]]

  **Quantile Data:**
  * 5th Percentile: [nFeature_quantiles["5%"]]
  * 95th Percentile: [nFeature_quantiles["95%"]]

  [Summary statistics for each quality metric]

  ** Total number of cells : ** [nCells]

  **Please provide the refined lower and upper cutoff values for nCount_RNA
and nFeature_RNA, [and the upper cutoff values for [quality_metrics]],
calculated based on the provided data and taking into account the potential
presence of tails in the distribution. Dont generate the cuffoff ONLY based
on the percentiles, instead combine the percentiles with a logic that we need
to remove outliers, but still keep good data. Make sure to state that the
researcher should test a range of values around your recommendation**. Do not
provide any IMPORTANT NOTES. Start the response saying that, based on the
your data summary, below are my recommendations for the quality filtering of
the data
  ```
```

Example SCassist Command and Output;

```
> SCassist_analyze_quality("KO", llm_server="google", api_key_file = api_key_file, model_G = "gemini-1.5-flash-latest")
```

Based on the provided data summary, below are my recommendations for the quality filtering of the single-cell RNA sequencing data:

**nCount_RNA:**

The distribution of `nCount_RNA` shows a substantial range, suggesting the presence of outliers. Simply using the 5th and 95th percentiles might be too restrictive, potentially removing valuable cells. To balance sensitivity and removal of outliers, I recommend the following:

* **Lower cutoff:** Instead of the 5th percentile (916.25), consider a value slightly higher, perhaps around 1500. This accounts for the lower tail but avoids being overly aggressive.

* **Upper cutoff:** The 95th percentile (27561.75) seems reasonable as an upper bound. However, consider exploring values slightly lower (e.g., 25000) to further mitigate the influence of potential high-count outliers.

**Recommended range for testing:** For `nCount_RNA`, test cutoffs in the ranges of 1000-2000 (lower) and 25000-30000 (upper).

**nFeature_RNA:**

Similar to `nCount_RNA`, the `nFeature_RNA` distribution exhibits a wide range. Again, a balance between outlier removal and data retention is crucial.

* **Lower cutoff:** A value slightly above the 5th percentile (511) is recommended. Consider a lower cutoff around 750.

* **Upper cutoff:** The 95th percentile (4897.5) appears suitable. Explore slightly lower values (e.g., 4500) to be more conservative.

**Recommended range for testing:** For `nFeature_RNA`, test cutoffs in the ranges of 600-800 (lower) and 4500-5000 (upper).

The researcher should test a range of values around these recommendations to optimize the filtering process and find the best balance between removing low-quality cells and retaining biologically relevant data. The optimal cutoffs will depend on the specific dataset and research question.
