## Supplementary File 3 for "SCassist: An AI Based Workflow Assistant for Single-Cell Analysis"

**Supplementary File 3:** Example Annotation Output from GPTCellType vs SCassist for the same FindAllMarkers input.

1. GPTCellType Command and Results

```
> gptcelltype(markersall, model = 'gpt-4o-mini')
```

[1] "Note: OpenAI API key found: returning the cell type annotations."

0 T cells

1 Natural Killer (NK) cells

2 Regulatory T cells (Tregs)

3 Mitochondrial cells (possibly representative of multiple cell types)

4 Cytotoxic T cells

5 Proliferating cells (e.g., cancer cells or activated lymphocytes)

6 Myeloid cells (e.g., macrophages or dendritic cells)

2. SCassist Command and Results

```
> sca_annotation <- SCassist_analyze_and_annotate(markersall,
top_genes = 10, seurat_object_name = "KO", llm_server="openai",
api_key_file = api_key_file_openai, model_C = "gpt-4o-mini")
```

[1] "The output of Analyze and Annotate is saved as a tab delimited text if your current working directory. If you provided the name of the seurat object, the annotation is also added to that object in the SCassist_annotation column"

| cluster_num | cell_type |
|---|---|
| 0 | NK cells |
| 1 | T cells |
| 2 | T regulatory cells |
| 3 | Mitochondrial cells |
| 4 | Cytotoxic T cells |
| 5 | Proliferating Cells |
| 6 | Macrophages |

scassist_reasoning

0 The expression of markers such as Klrd1 (also known as NKG2D), Klrk1 (also known as NKG2A), and Il2rb indicates a natural killer (NK) cell lineage, which is known for its role in the immune response against virus-infected and tumor cells. Additionally, Ahnak and Cst7 suggest involvement in cell signaling and cytotoxic function typical of NK cells, while the presence of Itgb1 (integrin beta 1) supports adhesion and navigation during immune responses.

1 The markers Cd3e, Lat2, and Trac indicate a T cell lineage, with Cd3e being a core component of the T cell receptor complex. The presence of Klrb1f and Klrc2 suggests potential involvement of natural killer T (NKT) cells or cytotoxic T cells, while Bcl11b is associated with T cell development. The overall marker profile strongly supports a T cell identity.

2 The presence of markers such as Foxp3, Ctla4, and Il2ra strongly indicates a population of T regulatory cells, which play a crucial role in maintaining immune tolerance

and preventing autoimmune responses. Additionally, the expression of Izumo1r and Tnfrsf4 suggests these cells may be involved in specialized interactions that modulate immune responses.

3 The presence of multiple mitochondrial genes such as mt-Nd2, mt-Nd4, and mt-Cytb, combined with ribosomal protein markers like Rpl37a, Rpl34, and Rps16, indicates a cell type with a high metabolic activity and significant mitochondrial function, typical of cells involved in energy production and respiration.

4 The presence of markers Cd8a and Cd8b1 strongly indicates that this cluster primarily consists of cytotoxic T lymphocytes, which are critical for immune response and pathogen clearance. Other markers such as Ly6c2 and Adgrg5 further support the role of these cells in immune functions, suggesting an activated state characteristic of effector T cells.

5 The presence of markers such as Birc5, Kif11, Cdca8, Ccna2, Cdk1, and Mki67 indicates a strong association with actively dividing cells, as these genes are known to be involved in cell cycle regulation and mitotic processes, suggesting that this cluster represents a population of rapidly proliferating cells, likely cancerous or stem cell-derived.

6 The presence of markers such as Sirpa, Fcgr4, and Csf1r suggests a myeloid lineage consistent with macrophage identity, indicating roles in phagocytosis and immune response. Additionally, markers like Adgre1 and Clec4a3 further support their involvement in tissue-resident or activated macrophage functions, highlighting their roles in inflammation and pathogen clearance.
