## Supplementary Data for "SCassist: An AI Based Workflow Assistant for Single-Cell Analysis"

### Supplementary Data in this File

**Use of Large Language Models:**

Authors declare that Google Gemini and Meta's Llama3 were used as an aid to correct written text, as an algorithmic technique for research study, as an evaluation technique to identify anomalies in our code and text, as a code writing assistant and for document creation. The authors have reviewed any content generated by Gemini and Llama3.

This supplementary file includes the following content:

Supplementary Table 3: Publicly available single-cell datasets used for SCassist evaluation.
Supplementary Table 4: Template scoring sheet (Likert scale) for Expert Human Evaluation.
Supplementary Table 5: Ground truth categories and Groundedness score.
Supplementary Table 6: BERT based semantic similarity scores for SCassist generated reports.
Supplementary Figure 1: Frequency Tables and Frequency Distribution of expert human evaluation scores.
Supplementary Table 7: Summary statistics on human expert evaluation, using the Likert Scale of 1 to 5. 1 being worst performing, 3 average performing and 5 best performing
Supplementary Table 8: The Wilcoxon Rank-Sum test showing no statistically significant differences in the LLM's performance across datasets and expert levels.

**Supplementary Table 3**: Publicly available single-cell datasets used for SCassist evaluation.

| Pubmed ID | Availability | Sample Type | Total No. of Cells |
|---|---|---|---|
| 39085199 | GitHub Link filtered_feature_bc_matrix.h5 | PBMCs from Birdshot Uveitis Patient | 14300 |
| 39085199 | GitHub Link filtered_feature_bc_matrix.h5 | PBMCs from Healthy Human Control | 24790 |
| 38697116 | NCBI GEO GSM6625298 Link filtered_feature_bc_matrix.h5 | NK, CD4+ and CD8+ T cells from LCMV infected WT mice | 5666 |
| 38697116 | NCBI GEO GSM6625299 Link filtered_feature_bc_matrix.h5 | NK, CD4+ and CD8+ T cells from LCMV infected Ifng - CTCF binding site mutant mice | 4885 |

**Supplementary Table 4**: Template scoring sheet (Likert scale) for Expert Human Evaluation.

| Score | Accuracy |
|---|---|
| 1 | The results and recommendations are completely inaccurate |
| 2 | The results and recommendations are slightly inaccurate |
| 3 | The results and recommendations are acceptable |
| 4 | The results and recommendations are mostly accurate |
| 5 | The results and recommendations are highly accurate |
| | **Relevance** |
| 1 | The results and recommendations are not relevant to this study at all |
| 2 | The results and recommendations are slightly relevant to this study |
| 3 | The results and recommendations are relevant to this study |
| 4 | The results and recommendations are mostly relevant to this study |
| 5 | The results and recommendations are highly relevant to this study |
| | **Clarity** |
| 1 | The results and recommendations are very difficult to understand |
| 2 | The results and recommendations are difficult to understand |
| 3 | The results and recommendations are clear |
| 4 | The results and recommendations are easy to understand |
| 5 | The results and recommendations are very easy to understand |
| | **Trustworthiness** |
| 1 | Doesn't make sense, so doesn't seem trustable at all |
| 2 | Doesn't make much sense, so doesn't seem very trustable |
| 3 | Makes some sense, so seems somewhat trustable |
| 4 | Makes sense, so seems trustable |
| 5 | Makes perfect sense, so seems completely trustable |
| | **Overall Satisfaction** |
| 1 | Completely useless, so would not recommend at all |
| 2 | Somewhat useless, would not recommend to most |
| 3 | Somewhat useful, would recommend with reservations |
| 4 | Useful, would recommend |
| 5 | Very useful, would highly recommend |

**Supplementary Table 5**: Ground truth categories and Groundedness score

| | Input Tokens | **SCassist-LCMV-Grounded** | Total reported | Groundedness score |
|---|---|---|---|---|
| Data Metrics | 16 | 14 | 14 | 100 |
| Variable Genes | 30 | 22 | 22 | 100 |
| Variance Explained | 10 | 10 | 11 | 90.91 |
| PC Genes | 250 | 73 | 73 | 100 |
| Annotation Genes | 630 | 234 | 234 | 100 |
| Enriched Terms | 3176 | 53 | 53 | 100 |
| Network Genes | 24 | 23 | 23 | 100 |
| **Average** | | | | **98.70** |
| **Total** | **4136** | | | |

| | Input Tokens | **SCassist-BCRUV-Grounded** | Total reported | Groundedness score |
|---|---|---|---|---|
| Data Metrics | 16 | 14 | 14 | 100 |
| Variable Genes | 30 | 29 | 29 | 100 |
| Variance Explained | 10 | 1 | 1 | 100 |
| PC Genes | 250 | 67 | 67 | 100 |
| Annotation Genes | 450 | 182 | 183 | 99.45 |
| Enriched Terms | 3327 | 28 | 28 | 100 |
| Network Genes | 27 | 25 | 25 | 100 |
| **Average** | | | | **99.92** |
| **Total** | **4110** | | | |

**Supplementary Table 6**: BERT based semantic similarity scores for SCassist generated reports

| | Input Tokens | **SCassist-LCMV-Similarity Score** |
|---|---|---|
| KEGG_pathways report | 99 | 0.76059038 |
| GO_terms report | 2196 | 0.73808815 |
| Network summary | 24 | 0.79054358 |
| **Total/Average** | **2319** | **0.763074037** |

| | Input Tokens | **SCassist-BCRUV-Similarity Score** |
|---|---|---|
| KEGG_pathways report | 117 | 0.7475318 |
| GO_terms report | 436 | 0.7515198 |
| Network summary | 27 | 0.7080501 |
| **Total/Average** | **580** | **0.735700567** |

**Supplementary Figure 1**: Frequency Tables and Frequency Distribution of expert human evaluation scores.

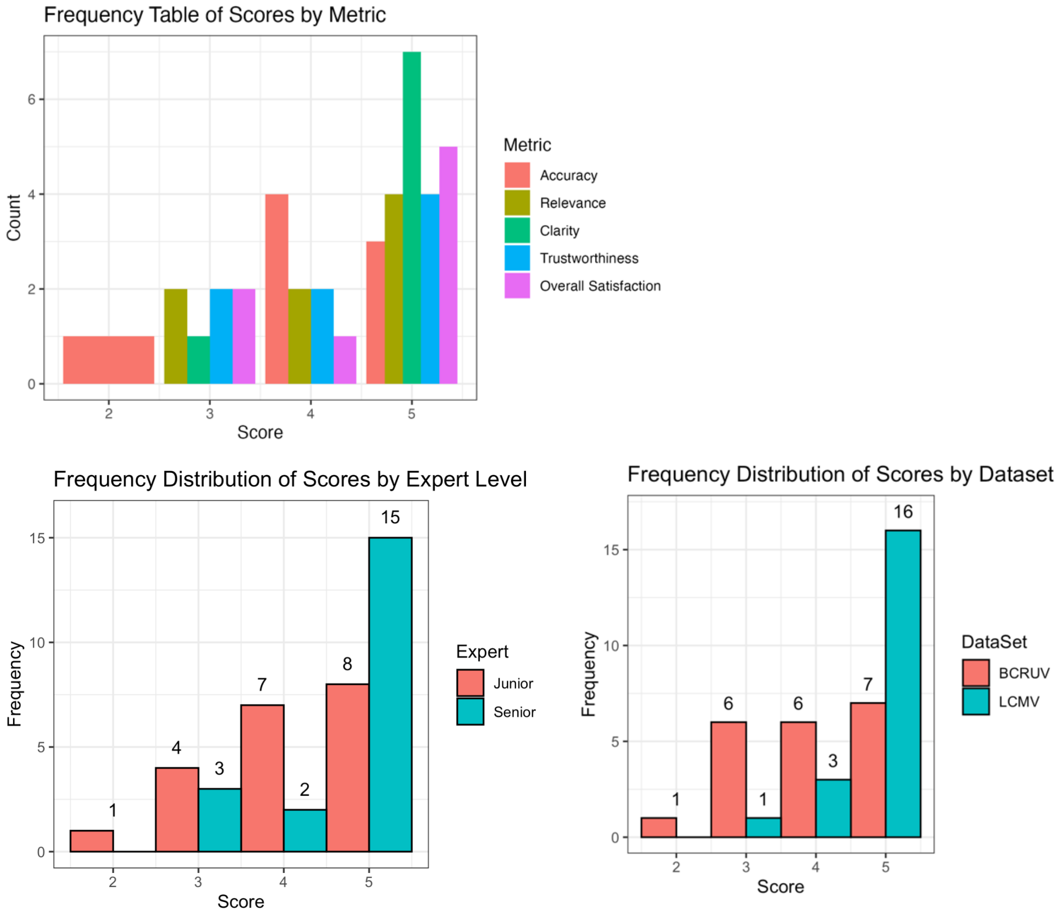

**Supplementary Table 7**: Summary statistics on human expert evaluation, using the Likert Scale of 1 to 5. 1 being worst performing, 3 average performing and 5 best performing

| Metric | Mean | Median | Min | Max | StdDev |
|---|---|---|---|---|---|
| **Accuracy** | 4.125 | 4 | 2 | 5 | 0.99 |
| **Relevance** | 4.25 | 4.5 | 3 | 5 | 0.89 |
| **Clarity** | 4.75 | 5 | 3 | 5 | 0.71 |
| **Trustworthiness** | 4.25 | 4.5 | 3 | 5 | 0.89 |
| **Overall Satisfaction** | 4.375 | 5 | 3 | 5 | 0.92 |

**Supplementary Table 8**: The Wilcoxon Rank-Sum test showing no statistically significant differences in the LLM's performance across datasets and expert levels.

| **BCRV vs LCMV** | |
|---|---|
| Metric | p_value |
| Accuracy | 1 |
| Relevance | 1 |
| Clarity | 1 |
| Trustworthiness | 0.80 |
| Overall Satisfaction | 0.33 |

| **Seniors vs Juniors** | |
|---|---|
| Metric | p_value |
| Accuracy | 0.29 |
| Relevance | 1 |
| Clarity | 1 |
| Trustworthiness | 1 |
| Overall Satisfaction | 1 |
